# Multiplexed biosensors for precision bacteria tropism *in vivo*

**DOI:** 10.1101/851311

**Authors:** Tiffany Chien, Tetsuhiro Harimoto, Benjamin Kepecs, Kelsey Gray, Courtney Coker, Kelly Pu, Tamjeed Azad, Tal Danino

**Affiliations:** Department of Biomedical Engineering, Columbia University, New York, NY 10027, USA; Data Science Institute, Columbia University, New York, NY 10027, USA; Herbert Irving Comprehensive Cancer Center, Columbia University, New York, NY 10027, USA

## Abstract

The engineering of microbes spurs biotechnological innovations, but requires control mechanisms to confine growth within defined environments for translation. Here we engineer bacterial growth tropism to sense and grow in response to specified oxygen, pH, and lactate signatures. Coupling biosensors to drive essential gene expression reveals engineered bacterial localization within upper or lower gastrointestinal tract. Multiplexing biosensors in an AND logic-gate architecture reduced bacterial off-target colonization *in vivo*.

## Main Text

Microbiome research continues to demonstrate the abundance of microorganisms in various environments (*1–3*). An emerging focus in synthetic biology engineers microbes to integrate within specific niches in nature, artificial infrastructure and the human body (*4–11*). Since genetic programming of bacteria enables them to sense and respond to physiological conditions *in situ*, this approach is poised to change existing paradigms for diagnosing and treating diseases such as inflammation (*12, 13*), infection (*14, 15*), and cancer (*16–18*). An essential element of this approach requires the precise regulation of microbial growth at disease sites, since uncontrolled bacterial replication can lead to severe side effects including tissue damage and septic shock *(19, 20)*. Therefore, engineering genetic circuits to control bacterial growth at specific regions provides a promising avenue to address this central challenge in translating next-generation microbial applications.

The majority of bacterial therapies thus far have relied on the natural tropism of bacteria in organs such as the gastrointestinal tract, skin, and tumors (*12, 17, 21–23*). While relying on inherent bacterial growth preference can control bacterial spatial localization, many bacteria can grow outside of their natural niche and can quickly spread to unintended locations. One approach to resolve this lack of specificity entails engineering approaches that controllably alter bacterial growth, including using metabolic auxotrophy, essential genes, toxins, and dependence on synthetic amino acids (*24–28*). Coupling such systems with an environmentally-responsive biosensor will contain engineered bacteria and prevent spreading to unintended tissues and targets (*29–31*), but precise targeting to specified organs remains a challenge. Here, we demonstrate programmable tropism and growth of bacteria within the body controlled by genetic circuits engineered to sense multiple distinct physiological signatures (**Fig. 1A**).

**Fig. 1.**
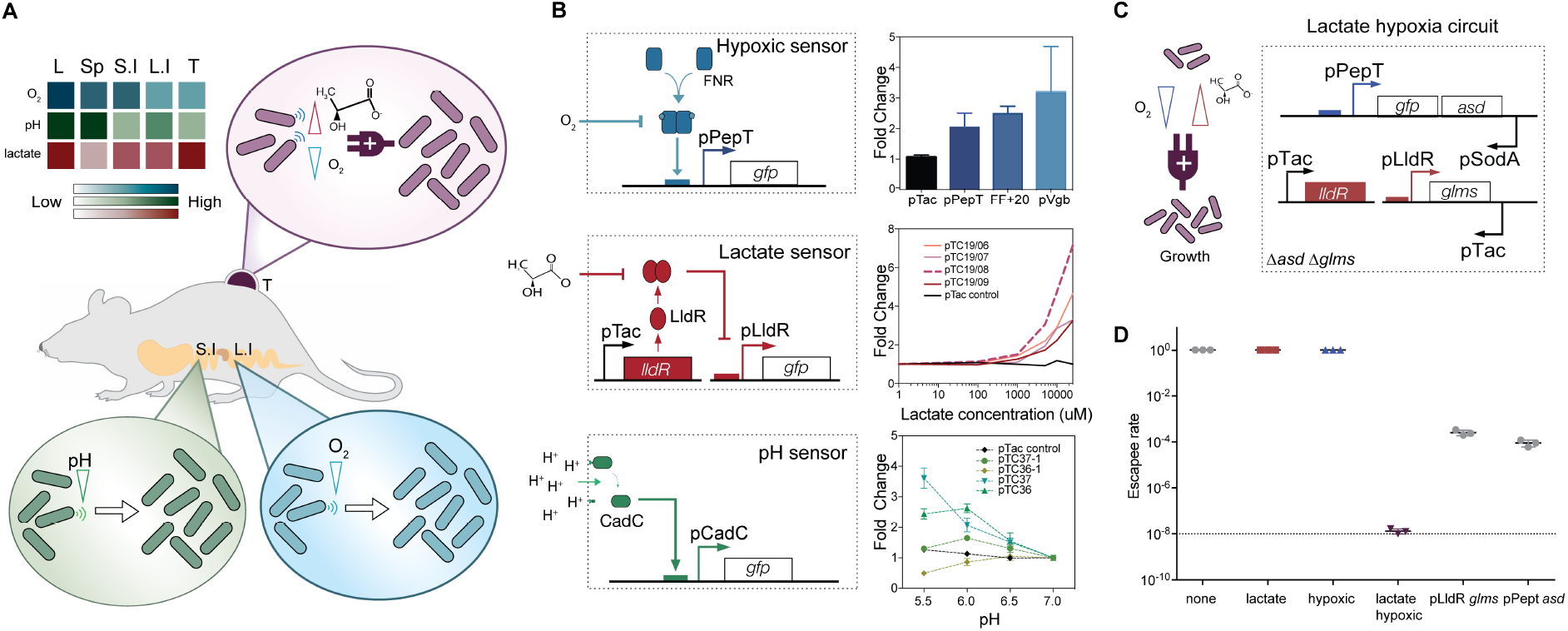
Design of biosensors for engineered bacteria tropism. **(A)** Engineered biosensors sense specific oxygen, lactate and pH levels in organs that enable precise growth tropism of bacteria *in vivo*. Different organs, ranging from Lungs (L), Spleen (Sp), Small Intestine (S.I.), Large Intestine (L.I.) and Tumor (T) exhibit varying bio-chemical signatures. **(B)** (Left) Each sensor architecture optimizes native bacteria promoters to sense specific environmental changes and express GFP. (Right) Bacteria were grown in a specified environment (0-20% oxygen, 0-10 mM lactate and pH of 5.5-7) for 16 hours, and fluorescence was measured (n=3, mean ±SEM). (**C**) Schematic of lactate-hypoxia AND logic gate circuit consists of both the hypoxic promoter pPepT driving essential gene *asd* and lactate sensor pLldR driving a second essential gene *glms*. (**D**) Escapee rates of lactate-hypoxia AND logic gate circuit were cultured in the different four conditions (none, 10mM lactate supplemented, hypoxic condition and combination of 10mM lactate with hypoxic condition). Two single circuits were grown in either lactate or hypoxic conditions (n=3, mean ±SEM). Samples were grown for 16 hours and then plated on LB agar supplemented with DAP and D-glucosamine. Colonies were counted after incubating at 37°C overnight. Dotted line indicates NIH guideline for engineered microorganism escapee rate.

To construct bacteria biosensors that can distinguish unique organ signatures, we chose oxygen, pH and lactate as physiological indicators, as they are predominantly found throughout the body (*32–36*). To sense oxygen, we utilized a hypoxia-sensing promoter (pPepT) that is primarily regulated by the transcriptional activator, fumarate and nitrate reduction regulatory protein (FNR) (*30, 37, 38*) (**Fig. 1B**). In the absence of oxygen, FNR binds a [4Fe–4S]^2+^ cluster to generate a transcriptionally active homodimer. However, the cluster is degraded in the presence of oxygen, which dissociates the FNR dimers into inactive monomers (*39*). Measuring GFP expressed under the control of the pPepT promoter on a plasmid, we detected elevated levels of fluorescence in response to hypoxic conditions (**Fig. 1B**). We next designed an L-lactate sensor, derived from the native LldPRD operon (*37*), to detect lactic acid fermentation by host cells. This system was constructed on two plasmids: a lactate-inducible reporter plasmid driving expression of a gene of interest and a repressor plasmid, which produces the repressor LldR that can dimerize to inhibit expression of the reporter gene unless bound to lactate (**Fig. 1B**). In response to increasing concentrations of L-lactate in the culture media, we observed a corresponding step-wise increase in GFP (**Fig. 1B**). Last, we engineered a pH sensitive promoter, pCadC, regulated by a membrane tethered activator protein (CadC) (*40–42*), which shows increased activity in acidic media compared to media at a neutral pH (**Fig. 1B**).

To tune the sensitivity level of biosensors with functional gradients in physiological conditions, we built a library of sensors. For hypoxic sensors, we utilized three distinct hypoxia promoters (pPepT, FF+20 and pVgb) (*30, 43, 44*) and demonstrated that hypoxia induces expression for each promoter compared to a constitutive control (**Fig. 1B**). For both lactate and pH sensors, we varied the copy numbers of the plasmids encoding the promoter and the regulatory proteins. Both sensors showed significant activation under increasing lactate concentrations and decreasing pH levels with varied sensitivity and dynamic ranges (**Fig. 1B, Fig. S1-2**). By tuning these parameters, variants demonstrated activation at physiological concentrations similar to naturally-occurring levels in various organs, such as the gastrointestinal tract and solid tumors (*45, 46*). Since bacteria are often subjected to multiple environmental conditions simultaneously *in vivo*, we also tested the cross-reactivity of the sensors under overlapping environmental conditions. We observed robust activity of sensors, regardless of the other conditions, which suggests the ability of using these sensors in an orthogonal manner (**Fig. S3**).

We next sought to restrict bacterial replication in response to distinct biochemical signatures by expressing key essential genes under the control of the biosensor promoters. We chose an essential gene (*asd*) involved in lysine, threonine, and methionine biosynthesis. This process can be rescued by adding diaminopimelic acid (DAP), a bacteria specific amino acid that can be supplemented in the media (*30, 47*). Importantly, DAP cannot be produced or metabolized from the host cellular environment, which provides an ideal strategy for biocontainment *in vivo*. We knocked out the *asd* gene from the genome of *E. coli* (**Fig. S4A**) and placed the gene under the control of the three biosensor promoters. Since the initial circuits showed high levels of basal expression, we reduced this expression by we constructed a library of variants with a range of gene copy numbers (colE1, p15a, or sc101 origin of replications and single genome integration) (**Fig. S5A, Table S1**). We measured the ratio of bacterial growth in non-permissive environments (no DAP in media) and permissive conditions (DAP in media), which represented the “escapee rate”for characterizing the containment capability of the engineered bacteria. We identified several strains with low escapee rates that could grow selectively under hypoxia, high lactate, or low pH levels (**Fig. S5B**). The proportions of bacteria that grew in non-permissive conditions (escapee rate) were less than 10^−4^, 10^−3^, 10^−2^ for hypoxia, lactate, and pH, respectively.

To further enhance the specificity of our biocontainment circuits, we designed an AND logic gate circuit, which only permits bacterial replication in the presence of two different environmental signatures. We engineered a strain with the lactate sensor driving *glms* and an additional gene orthogonal to *asd* that encodes for an essential glucosamine-6-phosphate synthase that can be rescued with D-glucosamine (*30, 47*). The engineered strain exhibited selective growth in the presence of 10 mM lactate with a 10^−3^ escapee rate (**Fig. S6**). By combining this system with a hypoxia-dependent *asd* auxotroph in a double knockout strain (**Fig. 1C**), we demonstrated growth only when both DAP and D-glucosamine supplements were added (**Fig. S4C**). When bacteria were cultured under both low oxygen and high lactate, the AND logic gate strain grew, while we measured no colonies up to the limit of detection (LOD) for hypoxic- or lactate-only culturing conditions (**Fig. 1D, Fig. S7**). The 10^−8^ escapee rate for the lactate hypoxic circuit improved containment 10^4^-fold compared to single containment modules alone and adheres to the NIH guideline for engineered microorganisms (**Fig. 1D**)(*48*). These findings highlight that using multiplexed sensor-based logic gate circuit architecture significantly improves growth specificity.

To progress toward the *in vivo* characterization of biosensors, we first assessed the capability of bacteria biosensors to sense metabolic activity of mammalian cell cultures *in vitro* (**Fig. 2A**). We cultured cell lines from various origins, including colorectal, lung, lymph nodes, and breast, over 5 days and measured the levels of lactate and pH in the culture media after collection (**Fig. S8**). Subsequently, we cultured sensor-containing bacteria in the collected cell media supernatant and measured the fluorescence from the bacteria sensor strains after 16 hours. We observed a concomitant increase in the bacterial fluorescent signal as lactate concentrations increased and the pH level decreased (**Fig. 2B, C**). To confirm that the promoter activity is independent of the cell type of origin, we assayed media supernatant samples from six different lung cancer cell lines and a lung fibroblast cell line. We found that the sensor showed an increase in fluorescent signal only in the cell lines that produced high levels of lactate (**Fig. S9**). To characterize the hypoxic sensor in culture media, we incubated the bacteria with cell media in a culture chamber with varying oxygen levels. We observed consistent increases in fluorescence signal following decreasing oxygen levels across cell lines (**Fig. 2D**). Collectively, these results demonstrate that our engineered bacteria can effectively sense biochemical signatures produced from host cellular metabolic activities.

**Fig. 2.**
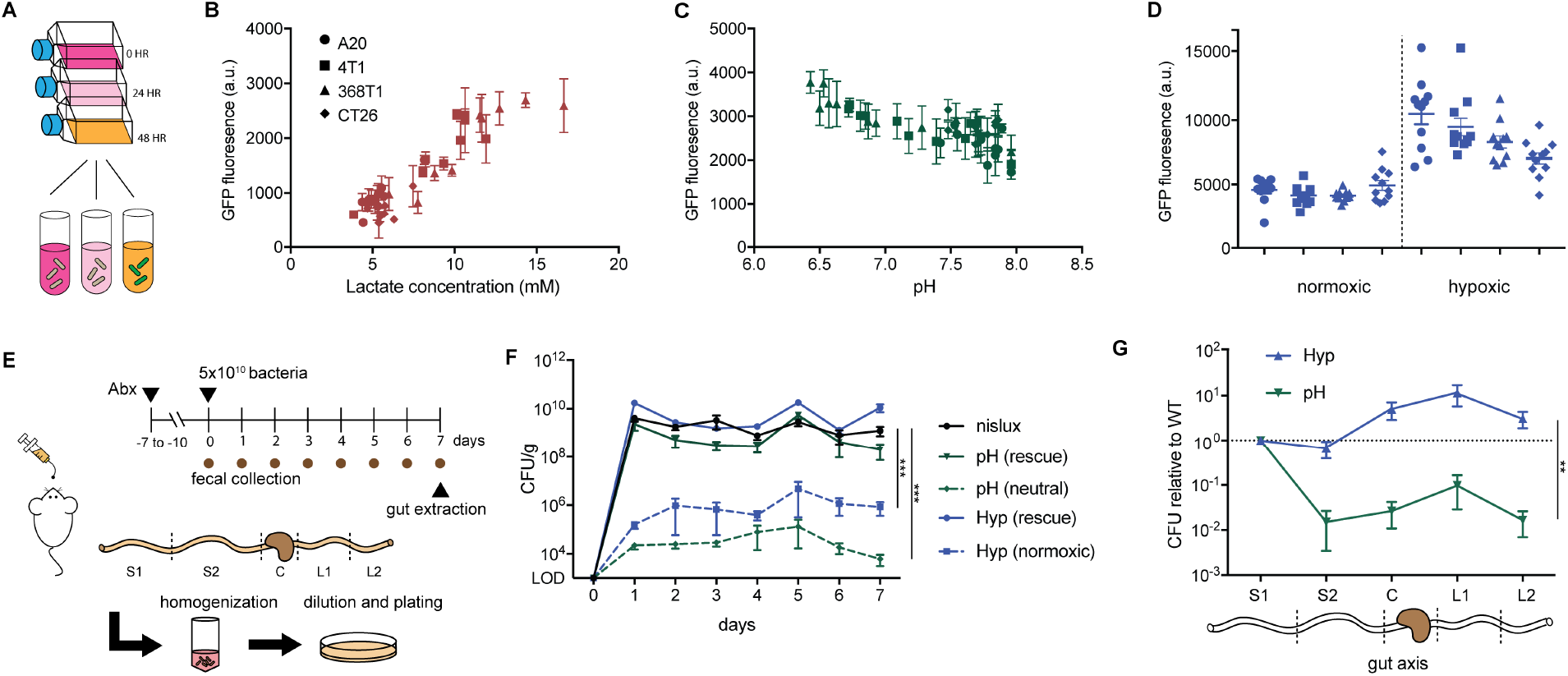
Engineered biosensors respond to physiological cues *in vitro* and enable redirected growth tropism and biocontainment in mouse gut. (**A)** Cell culture media supernatant from four cancer cell lines (A20, CT26, 368T1 and 4T1) were collected over five days and then cultured with our three bacteria sensors. (**B-D**) Fluorescence activation of lactate (**B**, red), pH (**C**, green), and hypoxic (**D**, blue) sensors when cultured in the collected supernatant overnight at 37°C. Hypoxic sensor cultured in cell media supernatant was grown in ±oxygen conditions. GFP signal was normalized by constitutive promoter control. (n=3, mean ±SEM). (**E**) Fresh fecal pellets were collected every day for 7 days, homogenized and plated on LB agar plates with antibiotic selection ±DAP. Mice were sacrificed at the end of the experiment (day 7), and the gastrointestinal tract was sectioned into 5 regions (S1, upper small intestine track; S2, lower small intestine track; C, caecum; L1, upper large intestine track; L2, lower large intestine track). The regions were homogenized and plated on LB agar plates with antibiotic selection and DAP. Colonies were counted the following day. (**F**) Colony counts from fresh fecal pellet homogenates were recovered by plating in permissive and non-permissive conditions (Nislux: n=7, pH: n=7, hypoxic: n=8 per time point, mean ±SEM. ***p = 0.0005 two-way analysis of variance (ANOVA) with Tukey’s multiple comparisons test). (**G**) Ratio of colonies for sensor containment circuit compared to wild type control along the gut axis. Data normalized to S1 region (Nislux: n=7, pH: n=7, hypoxic: n=8, mean ±SEM. **p = 0.007 two-way analysis of variance (ANOVA)).

Since oxygen and pH levels decrease along the longitudinal axis of the intestine (*46*), we exploited this system to engineer bacteria localization in the gut using our biosensor circuits. We transformed pH and oxygen sensors driving *asd* gene expression in a probiotic bacteria, *E. coli* Nissle 1917 (EcN), currently prescribed for oral administration in humans with gastrointestinal disorders (*49, 50*). Following oral delivery of bacteria, we collected fecal samples over several days to assess their ability to grow outside of the host environment (**Fig. 2E**). For both hypoxia and pH circuits, we grew approximately 100-fold less bacteria from the fecal samples over one week compared to wildtype EcN (**Fig. 2F**). Then, we rescued engineered bacteria using DAP supplementation in the agar to similar colony forming units (CFU) as in wildtype EcN. These results suggest that the reduced bacterial numbers outside of the host occurred through engineered auxotrophy. We then examined the bacterial distribution along the axis of the gut by measuring CFU from intestinal tissue (**Fig. 2E**). We found a 10-fold enrichment of hypoxia-dependent bacteria in the large intestine compared to the small intestine (**Fig. 2G**). In contrast, we observed a decrease in CFU toward the distal sites of the gastrointestinal tract for pH dependent bacteria (**Fig. 2G**). Together, these results demonstrate engineered bacterial growth tropism along the axis of the gastrointestinal tract based on physiochemical cues.

We next tested whether multiplexed containment circuits can enhance specificity for tumor environments. We employed a recently designed three-dimensional bacteria spheroid co-culture system (BSCC) that recapitulates properties of the tumor microenvironment including oxygen and nutrient gradients, mammalian cell metabolism, and local 3D growth of the bacterial population in tumors (*51*) (**Fig. 3A**). Attenuated *S. typhimurium*, commonly studied as a bacteria cancer therapy (*52*), was co-cultured. Once in the spheroid core, we observed an increase in the total fluorescence signal from bacteria carrying hypoxia, lactate, and pH sensors driving GFP (**Fig. 3B-D, Fig S10, 11**). We consistently measured the highest reporter signals in the center of the spheroid, reflecting the expected biochemical gradients in the spheroid core. The combined lactate-hypoxia AND logic gate containment circuit showed comparable tumor colonizing capabilities as wildtype bacteria in tumor spheroids (**Fig. 3E**). To determine whether this growth requires tumor microenvironment-specific cues, we co-cultured double knockout bacteria (*asd*-/*glms*-) containing a single sensor driving a single essential gene. Single sensor mutants could not effectively colonize tumor spheroids (**Fig. S12**), indicating that bacteria required both to facilitate growth.

**Fig. 3.**
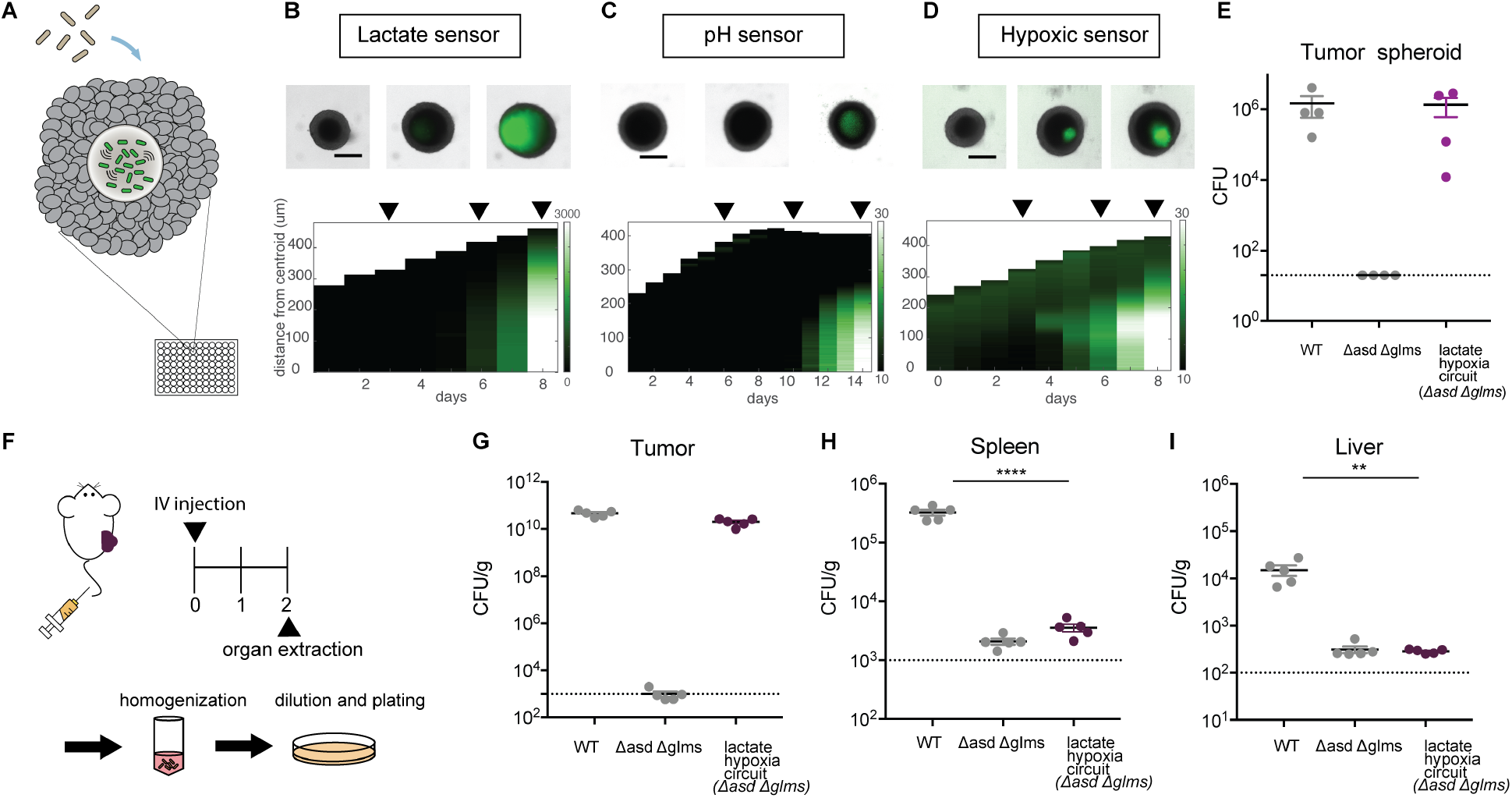
Multiplexed biosensor achieves enhanced specificity of bacteria tumor colonization. (**A**) Engineered bacteria biosensors were co-cultured in tumor spheroids and monitored for sensor activation. (**B-D**) (Top) Representative images of biosensors in tumor spheroids. (Scale bar, 200μm). (Bottom) Corresponding space-time diagram demonstrating radially averaged fluorescence intensity of lactate (**B**), pH (**C**) and hypoxia (**D**) sensors. The white boundary indicates the edge of the spheroid. (**E**) Recovered colony counts of wildtype *S. typhimurium* ELH1301, ELH1301 Δ*asd* Δ*glms* and hypoxia-lactate circuit tested in spheroid platform. All strains were co-cultured in spheroids for 6 days followed by homogenizing the spheroid and plating them on LB agar plates. (n=3, mean ±SEM). (**F**) BALB/c mice (n=5 per group) were implanted subcutaneously with 5 × 10^6^ CT26 cells on one hind flank. When tumor volumes were 100–150mm^3^, mice were intravenously administered ELH1301 (WT), double knockout (Δ*asd*Δ*glms*) or lacate-hypoxia circuit (in (Δ*asd*Δ*glms*). After 2 days, tumor, liver and spleen were homogenized and plated on LB agar plates with supplements. Bacteria colonizing tumor (**G**), spleen (**H**) and liver (**I**) tissues were quantified and counted after 1 day. Dotted line indicates the average limit of detection. (****p<0.0001, **p=0.0011 one-way analysis of variance (ANOVA) with Bonferroni’s multiple comparisons test).

We next intravenously injected our strains in a subcutaneous syngeneic mouse tumor model to mimic applications for bacteria cancer therapy (**Fig. 3F**). Organs and tumors were harvested and homogenized to assess bacterial colonization 2 days post-injection by CFU enumeration. As expected, we observed tumor colonization for both wildtype and lactate-hypoxia AND logic gate bacteria (**Fig. 3G**). These results confirm similar cancer targeting robustness between wildtype and engineered strains. Notably, low levels of the intravenously-introduced engineered bacteria were found in the spleen and liver (**Fig. 3H, I**) at levels similar to the auxotroph strains. In contrast, we recovered 10^4^ and 10^5^ CFU/g of wildtype bacteria from these organs respectively (**Fig. 3H, I**). Based on these results, we conclude that multiplexed bacteria decreased off-target colonization in the spleen and liver with a demonstrated >100-fold increase in tumor-targeting capability.

Our engineering proof-of-concept shown here demonstrates that bacteria can be engineered to grow locally only in distinct sections of the mouse gastrointestinal tract and tumors by utilizing specific distinct environmental signatures. By tuning and multiplexing biosensors responsive to physiological cues to construct biocontainment circuits, we demonstrated enhanced specificity and organotropic localization of microbial growth. This approach will enable precision targeting of specific physiological regions. We propose generalizing this approach to other bacteria species and organs for biomedical research and translation to human disease-states. As translation of engineered microbes to various technological applications undergoes continued implementation, robust and precise engineering of bacterial localization will provide a useful method to improve biocontainment and safety to target specific sites for local delivery of treatment options

## Acknowledgments

We thank J. Zhang for generating *glms* knock out strain and W. Mather for help with automated quantification of the BSCC platform.

## Author contributions

T.C., T.H. and T.D. conceived and designed the study. T.C., T.H., B.K., T.A. performed in vitro characterization of biosensor. T.C., T.H., K.G., C.C., K.P. performed in vivo experiments for this study. T.C., T.H. and T.D. wrote the manuscript.

## Data and materials availability

All data is available in the main text or the supplementary materials.

